# P-glycoprotein detoxification by the Malpighian tubules of the desert locust

**DOI:** 10.1101/529750

**Authors:** Marta Rossi, Davide De Battisti, Jeremy E. Niven

**Author notes:** Corresponding authors (MR) and (JEN).

## Abstract

Detoxification is essential for allowing animals to remove toxic substances present in their diet or generated as a biproduct of their metabolism. By transporting a wide range of potentially noxious substrates, active transporters of the ABC transporter family play an important role in detoxification. One such class of transporters are the multidrug resistance P-glycoprotein transporters. Here, we investigated P-glycoprotein transport in the Malpighian tubules of the desert locust (*Schistocerca gregaria*), a species whose diet includes plants that contain toxic secondary metabolites. To this end, we studied transporter physiology using a modified Ramsay assay in which *ex vivo* Malpighian tubules are incubated in different solutions containing the P-glycoprotein substrate dye rhodamine B in combination with different concentrations of the P-glycoprotein inhibitor verapamil. To determine the quantity of the P-glycoprotein substrate extruded we developed a simple and cheap method as an alternative to liquid chromatography–mass spectrometry, radiolabelled alkaloids or confocal microscopy. Our evidence shows that: (i) the Malpighian tubules contain a P-glycoprotein; (ii) tubule surface area is positively correlated with the tubule fluid secretion rate; and (iii) as the fluid secretion rate increases so too does the net extrusion of rhodamine B. We were able to quantify precisely the relationships between the fluid secretion, surface area, and net extrusion. We interpret these results in the context of the life history and foraging ecology of desert locusts. We argue that P-glycoproteins play an important role in the detoxification by contributing to the removal of xenobiotic substances from the haemolymph, thereby enabling gregarious desert locusts to maintain toxicity through the ingestion of toxic plants without suffering the deleterious effects themselves.

## Introduction

Insect excretory systems consist primarily of the Malpighian tubules and the hindgut, which act synergistically to regulate haemolymph composition [1,2]. Malpighian tubules are blind ended tubules that float in the haemolymph and empty into the gut at the midgut-hindgut junction, secreting primary urine, the composition of which is modified by water and ion reabsorption in the hindgut [3]. The tubules are considered analogous to vertebrate nephrons [2]. Cells of the epithelium forming the tubule wall express primary and secondary active transporters that move K^+^, Na^+^ and Cl^-^ ions into the lumen creating an osmotic gradient that produces water secretion (for a review see [4]). Insects regulate ion and water secretion according to their feeding habits and ecological niche. For example, haematophagous insects must cope with an excess of NaCl and water after a blood meal [5], whereas phytophagous insects must often cope with a diet rich in K^+^ as well as with secondary metabolites [6,7].

In addition to osmoregulation, Malpighian tubules play a fundamental role in the removal of metabolic waste and potentially noxious substances that have been ingested [1,8]. Alkaloids and organic anions and cations are actively transported by ATP-dependant transporters such as the multidrug resistance-associated protein 2 (MRP2) and P-glycoproteins (P-gps, multidrug resistance protein (MDR1) or ABCB1), both members of the ABC transporter family [9,10]. Multidrug resistance-associated protein 2 (MRP2) transporters are involved in the transport of organic anions [11,12], while P-glycoproteins transport type II organic cations (>500 Da), hydrophobic and often polyvalent compounds (e.g. alkaloids and quinones) [13].

The presence and physiology of these multidrug transporters have been explored using specific substrates and selective inhibitors (e.g. [9,11,14]). In the Malpighian tubules of the cricket (*Teleogryllus commodus*) and the fruit fly (*Drosophila melanogaster*), the transepithelial transport of the fluorescent MRP2 substrate Texas Red is reduced by the MRP2 inhibitors MK571 and probenecid [11], while the transport of the fluorescent P-glycoprotein substrate daunorubicin is selectively reduced by the P-glycoprotein inhibitor verapamil [11]. The transport of nicotine by P-glycoprotein transporters has also been demonstrated in numerous insect species, including the tobacco hornworm (*Manduca sexta* [9]), fruit fly (*D. melanogaster*), kissing bug (*Rhodnius prolixus*), large milkweed bug (*Oncopeltus fasciatus*), yellow fever mosquito (*Aedes aegypti*), house cricket (*Acheta domesticus*), migratory locust (*Locusta migratoria*), mealworm beetle (*Tenebrio molitor*), American cockroach (*Periplaneta americana)* and cabbage looper (*Trichoplusia ni)* [15]. In insects, the understanding of the interaction between xenobiotics (i.e. insecticides, herbicides, miticides and secondary plant metabolites) and P-glycoprotein transporters is still limited, but there is an increasing interest in understanding how different xenobiotics can act synergistically to maximize the efficacy of insecticides in pests or impair the xenobiotic detoxification of beneficial insects such as honey bees [16].

Desert locusts (*Schistocerca gregaria*) are generalist phytophagous insects with aposematic coloration in the gregarious phase. They feed on a wide range of plants including those, such as *Schouwia purpurea* and *Hyoscyamus muticus*, that contain toxins to become unpalatable and toxic to predators [17–20] Nevertheless, it is likely that gregarious desert locusts excrete some of the toxins that they ingest, relying instead on their gut contents to maintain toxicity [21,22]. Two lines of evidence suggest that this excretion is likely to involve P-glycoproteins: (1) they are expressed in the Malpighian tubules of numerous species (e.g. *A. domesticus, L. migratoria, P. americana*) from orthopteroid orders [15]; and (2) they are expressed in the blood brain barrier of the desert locust [23]. However, P-glycoproteins in the Malpighian tubules of desert locusts have not been studied previously.

Here we show that xenobiotic transport and extrusion in the Malpighian tubules of the desert locust is an active process dependent upon P-glycoprotein like transporters using isolated tubules to perform a modified Ramsay secretion assay [24]. We evaluated the extrusion of the P-glycoprotein substrate dye rhodamine B (e.g. [25,26]) with or without the addition of the selective P-glycoprotein inhibitor verapamil (e.g. [23,27,28]). Our results suggest that P-glycoprotein transporters may play an important role in the xenobiotic detoxification in the Malpighian tubules of the desert locust. By using linear mixed effect models to account for repeated observations of single tubules and obtaining multiple tubules from single locusts, we found that tubule surface area more accurately predicts fluid secretion rate than diameter or length. Moreover, this statistical model allowed us to quantify the influence of the surface area on the fluid secretion rate in different treatments, and how it changes over time. We found that the surface area of the tubules positively influences their fluid secretion rate and that the fluid secretion rate influences the net extrusion of rhodamine B. We propose that this assay may be used in future to understand the physiology of the P-glycoproteins when exposed to a wide range of different substances.

## Materials and methods

### Animals

Fifth instar desert locusts (*Schistocerca gregaria;* Forskål, 1775) were obtained from Peregrine Livefoods (Essex, UK) and raised under crowded conditions at 28-30°C with 12:12 photoperiod. Locusts were fed with organic lettuce, fresh wheat seedlings and wheat germ *ad libitum*. Fifth instar nymphs were checked daily and, within 24 hours post-eclosion, were marked with acrylic paint (Quay Imports Ltd, Kirkham, Lancashire, UK). Only adult males between 20 and 22 days post-eclosion were used in the experiments.

### Saline and chemicals

The saline used was adapted from the Ringer solution of Maddrell and Klunsuwan [6]. Its composition was: 5.73 g/L NaCl (98 mM), 0.30 g/L KCl (4 mM), 2 mL CaCl_2_ solution 1M (2 mM), 1.86 g/L NaHCO_3_ (22 mM), 1.09 g/L NaH_2_P0_4_·2H_2_O (7 mM), 0.19 g/L MgCl_2_ (2 mM), 1.80 g/L glucose (10 mM), 0.83 g/L sodium glutamate (4.9 mM), 0.88 g/L sodium citrate (3.5 mM) and 0.37 g/L malic acid (2.8 mM). The final pH of the saline was 7.15. It was stored at 4°C for a maximum of three days. Stock solutions of rhodamine B (50 mM and 3 mM) and verapamil hydrochloride (20 mM) were prepared in water and diluted to the final concentration in the saline. Rhodamine B was applied at 60 μM and verapamil at 125 μM or 250 μM. All chemicals were purchased from Sigma-Aldrich (UK) or Fisher Scientific (UK).

### Locust dissection

The locusts were placed in the freezer for 4-5 minutes until sedated. Upon removal from the freezer, the abdomen was cut transversely ~5 mm from the anus, and holding the head with one hand and the thorax with the other hand, the head was pulled away from the remainder of the body (Fig 1A) [6]. In this way, the entire gut with the Malpighian tubules attached was removed from the body. The gut was placed onto an 8 cm Sylgard^®^ 184 (Dow Corning, Midland, MI, USA) coated Petri dish, filled with saline. The head was separated from the saline using modelling clay (Plasticine^®^) (Fig 1B). The preparation was pinned at the cut distal end of the gut to prevent it from floating.

**Fig 1.**
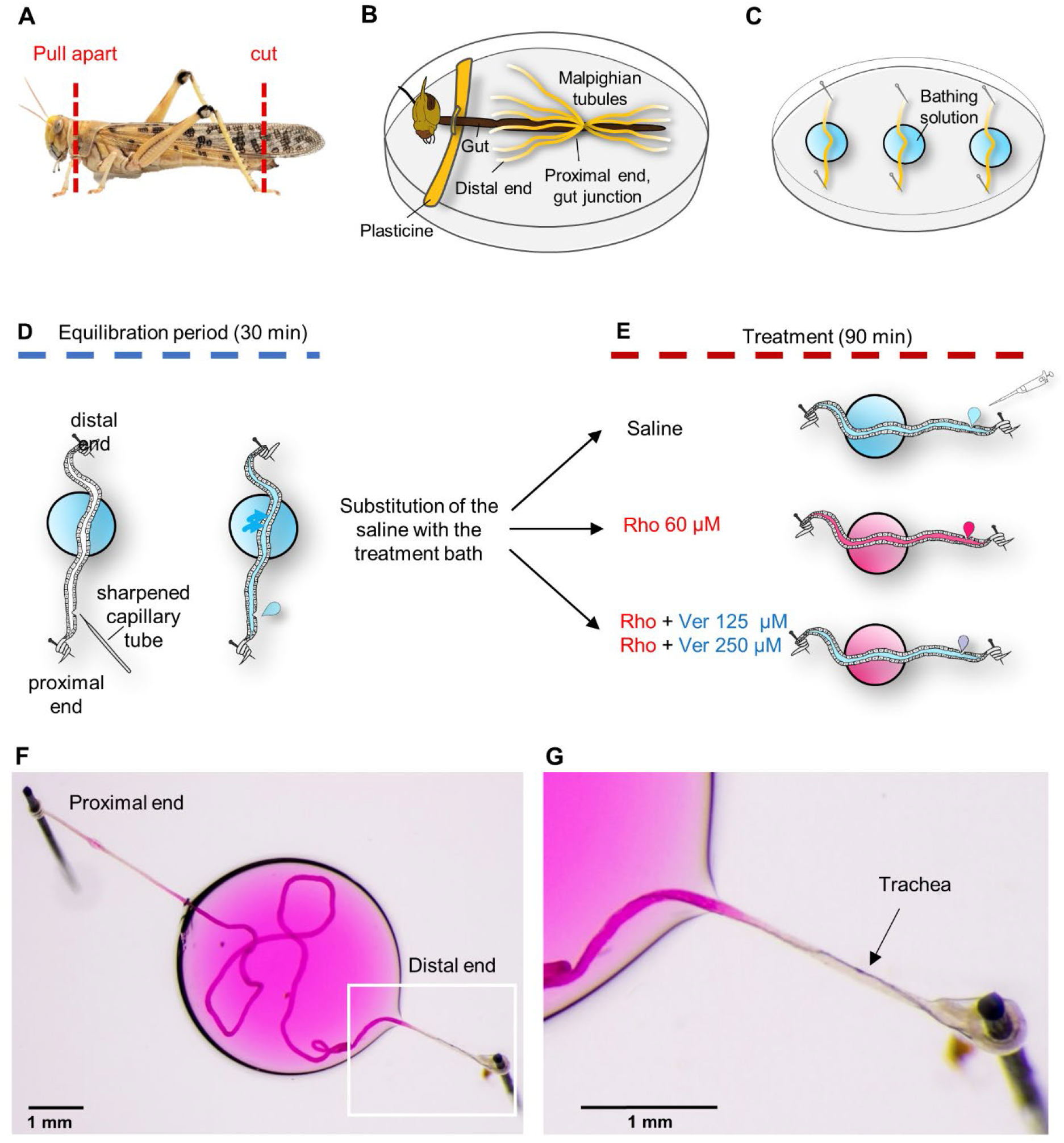
Preparation of Malpighian tubules for *ex vivo* experimentation from the desert locust, *Schistocerca gregaria*, and experimental scheme for assaying the presence of P-glycoproteins within the Malpighian tubules. (**A**) Cutting the posterior tip of the abdomen permits removal of the head and the fully intact gut. (**B**) The gut is submerged in saline and the Malpighian tubules are exposed. The head is separated from the saline bath by a barrier of modelling clay. (**C**) Three tubules are removed from each locust and fixed on a Sylgard^®^ surface traversing a small drop of bathing solution, and covered with paraffin oil. (**D**) The proximal and distal ends of individual tubules are wound around Minuten pins to fix them. Once the saline bath and paraffin oil have been applied, the proximal end of the tubule is punctured with a sharpened capillary tube to allow fluid secretion. After 30 minutes of equilibration the saline bath was replaced by a bath containing one of the treatments. (**E**) Every 30 minutes the secreted droplet was removed, placed on the Petri dish and photographed. (**F**) An Example of an isolated Malpighian tubule to perform a modified Ramsay secretion assay. The middle section of the tubule is immersed in the bathing solution with the respective treatment, while the proximal and distal ends are fixed outside. (**G**) Detail of the distal end with the trachea visible. Only a small part of the trachea is immersed in the bath.

### Malpighian tubules dissection

Using a Nikon SMZ-U (Nikon Corp., Tokyo, Japan) stereoscopic microscope, the Malpighian tubules were removed by gently pulling the distal part to release them from the gut and cutting the proximal end at ~5 mm from the gut). Each isolated tubule was moved immediately into a 30 μL drop of saline on a 5 cm Sylgard^®^ coated Petri dish and covered with paraffin oil to prevent desiccation. Both ends of each Malpighian tubule were pulled out from the saline drop in opposite directions and wrapped around steel pins pushed into the Sylgard^®^ layer (Fig 1C,D). Three anterior tubules were removed from each locust. Tracheae coiled around the distal part of the tubule were not removed to prevent any damage of the tubule surface (Fig 1F,G).

### Fluid secretion (Ramsay) assay

Using a sharpened glass capillary tube, each tubule was punctured near the proximal end to allow the fluid secretion (Fig 1D). The tubule was allowed to equilibrate for 30 minutes at which point the saline bath was replaced with 30 μL drop containing one of the four different treatments we tested: (i) saline, (ii) rhodamine B 60 μM, (iii) rhodamine B 60 μM + verapamil 125 μM, (iv) rhodamine B 60 μM + verapamil 250 μM (Fig 1E). The first droplet secreted after the bath replacement was discarded. For the subsequent 90 minutes, the secreted droplet was removed at intervals of 30 minutes (Fig 1E,F) using a P10 pipette (Gilson Scientific UK, Dunstable, Bedfordshire, UK) and photographed with a digital camera (Canon EOS 7D; Canon, Tokyo, Japan) mounted with two custom attachments (Best scientific A clamp via 1.6 x Canon mount; Leica 10445930 1.0 x) on the stereoscopic microscope (Nikon SMZ-U; Nikon Corp., Tokyo, Japan). Images were shot in raw format and processed with ImageJ v.1.51p software [29]. To prevent the photobleaching of the rhodamine B, we minimised light exposure by conducting the experiment under red light and keeping the sample in a custom designed dark box between measurements.

### Droplet measurement

The diameter (μm) of each secreted droplet (S1 Fig) was measured to calculate its volume (nL) using the sphere formula, where *V* is the drop volume and *d* the droplet diameter. The volume was converted from μm^3^ to nL using the formula: 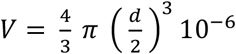.

For each tubule, we calculated the fluid secretion rate (nL/min), given by the droplet volume divided by the time between samples (30 mins). For each droplet, we also measured colour intensity to estimate the rhodamine B concentration (μM) from a calibration curve (see below). To estimate the number of moles of rhodamine B extruded per minute, we calculated the net extrusion of rhodamine B (fmol/min) as the product of the fluid secretion rate and the rhodamine B concentration.

### Rhodamine B calibration curve

The intensity of the droplets depends not only on rhodamine B concentration, but also on the droplet diameter (S2 and S3 Figs). So, we constructed a calibration curve for rhodamine B concentration to estimate the rhodamine B concentration of the droplets secreted. We prepared standard solutions of known rhodamine B concentrations: 0, 15, 30, 50, 60, 75, 120, 150, 240 and 480 μM. For each concentration, droplets of different sizes were placed on a Petri dish coated with Sylgard^®^ and covered with paraffin oil. We photographed the droplets against a white background at the same light intensity, and white balancing the camera before shooting. All the images were analysed subsequently using ImageJ v.1.51p software [29].

Droplet colour varied from white (transparent droplet at rhodamine B concentration = 0 μM) to intense pink, depending upon the rhodamine B concentration. We split each image into the component colour channels and measured the intensity of the green channel. To control for the background, we compared the mean intensity inside the droplet with that outside using the formula *I* = *I_i_ − I_o_*, where *I* is the intensity, *I_i_* is the intensity inside the droplet, and *I_o_* is the intensity outside the droplet. We used a range of droplet diameters from 138 μm to 999 μm.

To validate the reliability of using the green channel, we also measured the magenta channel and the total intensity using Adobe Photoshop CC v. 19.1.1 (Adobe Systems Incorporated, CA, USA). Both the magenta channel and total intensity correlated with the intensity of green channel (Pearson’s correlation, Magenta: p<0.001, df=17, R^2^=1; Total intensity: p<0.001, df=17, R^2^=0.99).

The relationship between the intensity and the rhodamine concentration depends upon the diameter of the droplet (S2 Fig). For each droplet diameter rank, the relationship between the intensity measured and the known rhodamine concentration can be described by a linear model. To determine the rhodamine concentration given the intensity and the diameter of the droplets, for each diameter rank we ran a linear regression model forced through the origin, with intensity as the independent variable and rhodamine concentration as the response variable (linear model: rhodamine concentration ~ intensity – 1). Hence, for each diameter rank we obtained the equation that predicts the rhodamine concentration from the intensity measured, given a specific diameter (S2 Fig).

The slope of the linear equations decreases as the diameter increases, following an exponential decay (S3A Fig). To obtain the equation that predicts the slope of the linear equations for a given diameter, we log transformed both axis and we ran a log-level regression model (Linear model: log (slope) ~ log (droplet diameter); S3B Fig). The resulting equation was: log(*slope*)_*i*_ = −1.44 · log(*diameter*)_*i*_ + 10.38.

Using the predict command in R, we used the model to predict the slope value (back transformed to the original scale) for each diameter of the droplets we collected in the experiment. Finally, we multiplied the slope value for the intensity measured to calculate the rhodamine concentration of each droplet.

### Malpighian tubule measurement

At the end of the assay, the tubule was photographed to measure its diameter (μm). The length (mm) of the tubule in contact with the treatment solution, was measured by cutting off the two extremities of the tubule outside the bath, laying the remaining section of tubule flat on the Sylgard^®^ base, and photographing it. The surface (mm^2^) of the tubule in contact with the bath, was calculated from the cylinder formula: 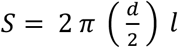, where *S* is the tubule surface, *d* is the tubule diameter, and *l* is the tubule length.

### Statistical Analysis

All the statistical analysis was conducted in R version 3.4.1 [30]. We performed Linear Mixed Effect Models (LMEM) by restricted maximum likelihood (REML) estimation by using the lmer function from the ‘lme4’ package [31]. We used the Akaike information criterion (AIC) [32] for model selection. Significances of the fixed effects were determined using Satterthwaite’s method for estimation of degrees of freedom by using the anova function from the ‘lmerTest’ [33]. The non-significant interactions (P>0.05) were removed. However, we retained all the main effects even if they were not statistically significant to avoid an increase in the type I error rate [34]. Estimated marginal means and pairwise comparisons were obtained using the ‘lsmeans’ package [35] and the p value adjusted with the Tukey method. All plots were made using the ‘ggplot2’ package [36].

To investigate the effect of the treatments on the fluid secretion rate and the net extrusion of rhodamine B, we analysed the interaction between the treatment (categorical), time of incubation (categorical) and the surface area (continuous (mm^2^)). For the rhodamine B concentration, we analysed the interaction between treatment (categorical) and time (categorical). To account for the nested structure of data, we included the individual locust as random intercept in the model. We also included tubule identity as a random intercept and time as random slope to account for the repeated measurements on the same individual tubule. To investigate the effect of the fluid secretion rate on the net extrusion of rhodamine B we analysed the interaction of the variables secretion rate, treatment and time including as before the individual locust as random intercept, and tubule identity as a random intercept and time as random slope. To simplify the interpretation of the regression estimates, we centred the surface variable on its mean. Therefore, all the estimates and comparisons are referred to a tubule with a mean surface area.

## Results

We prepared three Malpighian tubules from each locust (see Materials and Methods; Fig 1). Each tubule was punctured near the proximal end to allow the luminal fluid to be secreted and then they were allowed to equilibrate in the saline bath for 30 minutes (Fig 1D). The saline bath was then replaced with one of four treatments: saline (control); rhodamine B 60 μM (R60); rhodamine B 60 μM + verapamil 125 μM (V125); and rhodamine B 60 μM + verapamil 250 μM (V250) (Fig 1E). Six locusts were used for each of the treatments except for the R60 treatment in which eight locusts were used. Every 30 minutes the droplet secreted by the tubule during the Ramsay assay was removed.

### Fluid secretion rate and surface area

We determined the fluid secretion rate of each tubule from the volume of the droplet secreted after each 30-minute interval up to 90 minutes after the start of the treatment. Thus, for each tubule we had three measurements of the secretion rate in each of the four treatments. In total, there were 233 treatment observations (one droplet was lost after 60 minutes for the V250 treatment) from 78 Malpighian tubules.

To determine whether the surface area of the Malpighian tubules exposed to the bath solution influences the fluid secretion rate, we measured the length and diameter of each tubule immersed in the saline or treatment. By comparing linear mixed effect models that incorporated these measurements of length, diameter or surface area, we determined that surface area was the best explanatory variable (S1 Table). There was no difference in the surface area of Malpighian tubules exposed to the bathing solution among the treatments (F_3,22.02_=0.488, p=0.694; Control: 2.00 ± 0.09 mm^2^ (mean ± S.E.); R60: 2.01 ± 0.06 mm^2^; V125: 2.27 ± 0.06 mm^2^; V250: 2.29 ± 0.07 mm^2^).

The surface area of the tubule exposed in the bathing solution influenced the fluid secretion rate depending on the treatment (F_3,66.29_=3.25, p=0.027; Fig 2; Table 1A). Throughout the whole period of incubation, the surface area positively influenced the fluid secretion of tubules incubated in R60, V125 and Saline, while the V250 treatment the tubules showed no significant correlation between surface area and fluid secretion rate (Fig 2C; Table 1A). Having incorporated tubule surface area into our statistical model, we were able to compare the fluid secretion rates of our control and treatments objectively. The fluid secretion rate decreased over time irrespective of the treatment (F_2,82.36_=46.12, p=<.001; Time 60 vs Time 30: −0.12 ± 0.01 nL/min, p<.001; Time 90 vs Time 60: −0.04 ± 0.01 nL/min, p=0.013; Fig 3, Table 1B) and at each time point there was no significant difference between treatments (Fig 3, Table 1C).

**Table 1.**
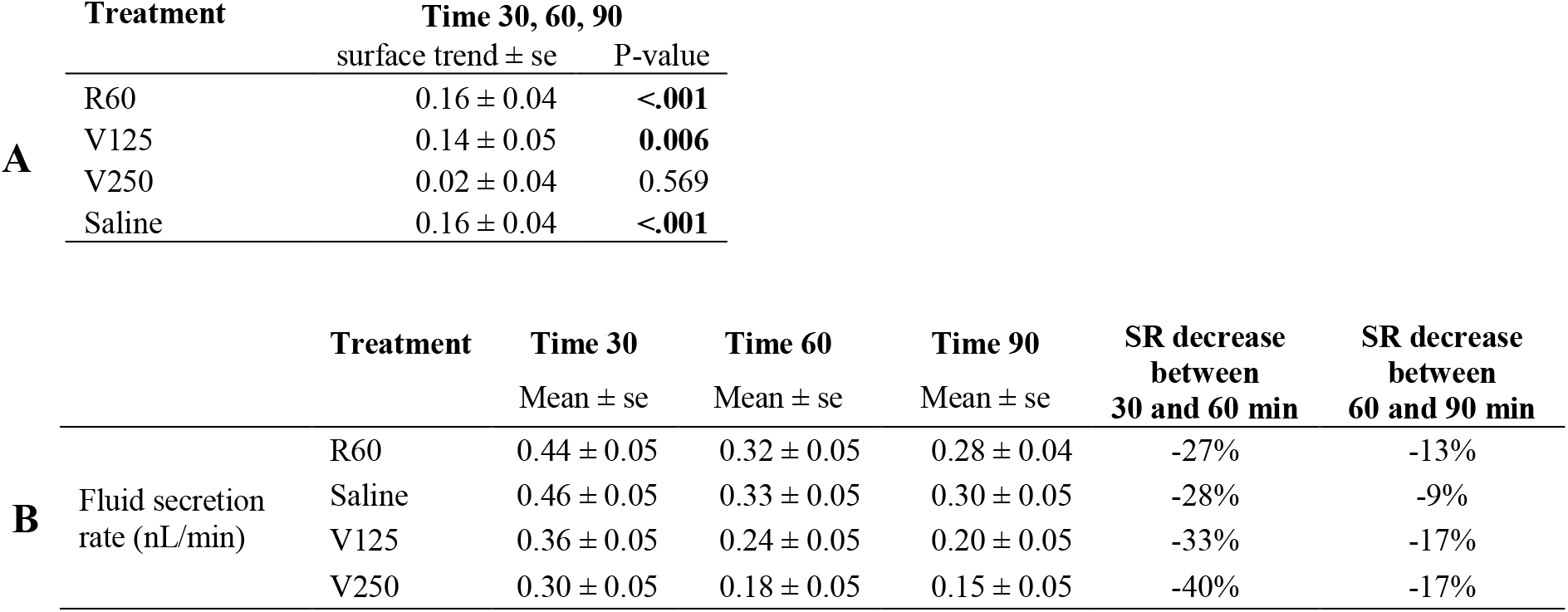
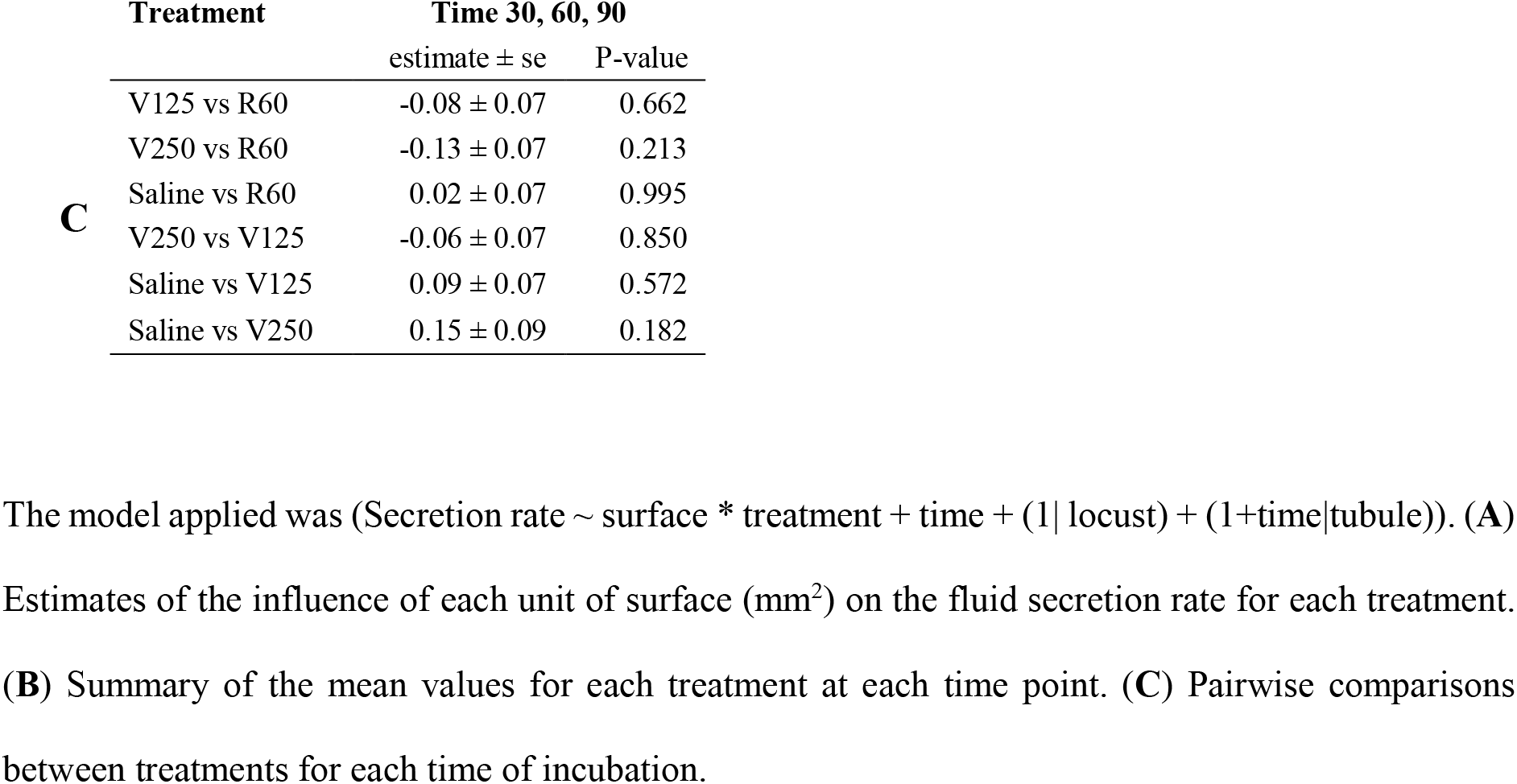
Outcomes of the linear mixed effect model investigating the effect of time of incubation, treatment, and surface area on the fluid secretion rate (SR) of Malpighian tubules.

**Fig 2.**
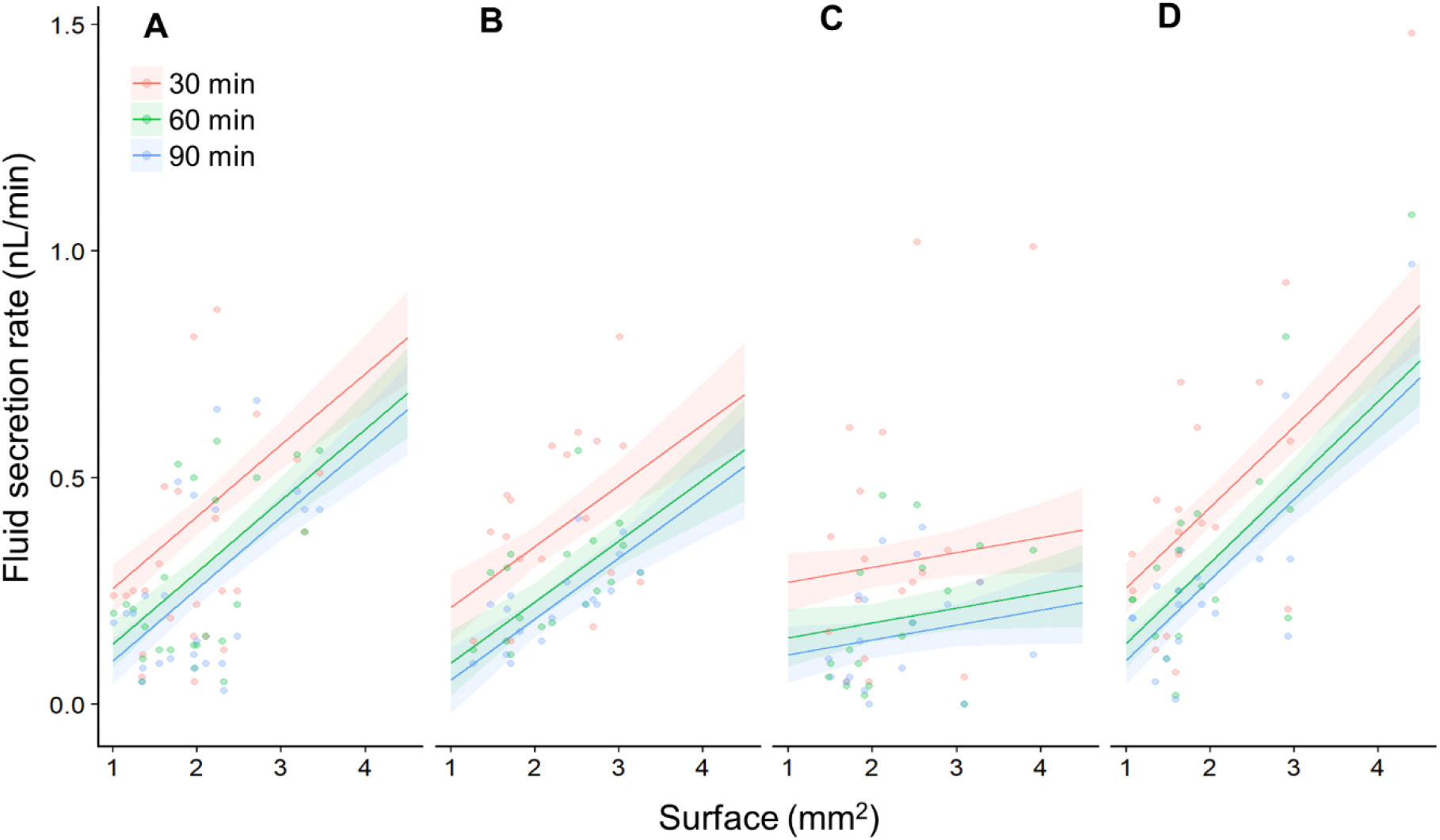
Tubule surface area positively influences the fluid secretion rate. (**A**) A plot of the surface area of the tubule exposed to the bath versus the fluid secretion rate for the R60 treatment every 30 minutes. (**B**) As in ‘A’ but for the V125 treatment. (**C**) As in ‘A’ but for V250 treatment. (**D**) As in ‘A’ but for the saline treatment. Small circles indicate the fluid secretion rates of individual tubules at a particular time point. Each line with the shaded area around represents the mixed effect model fit with the standard error.

**Fig 3.**
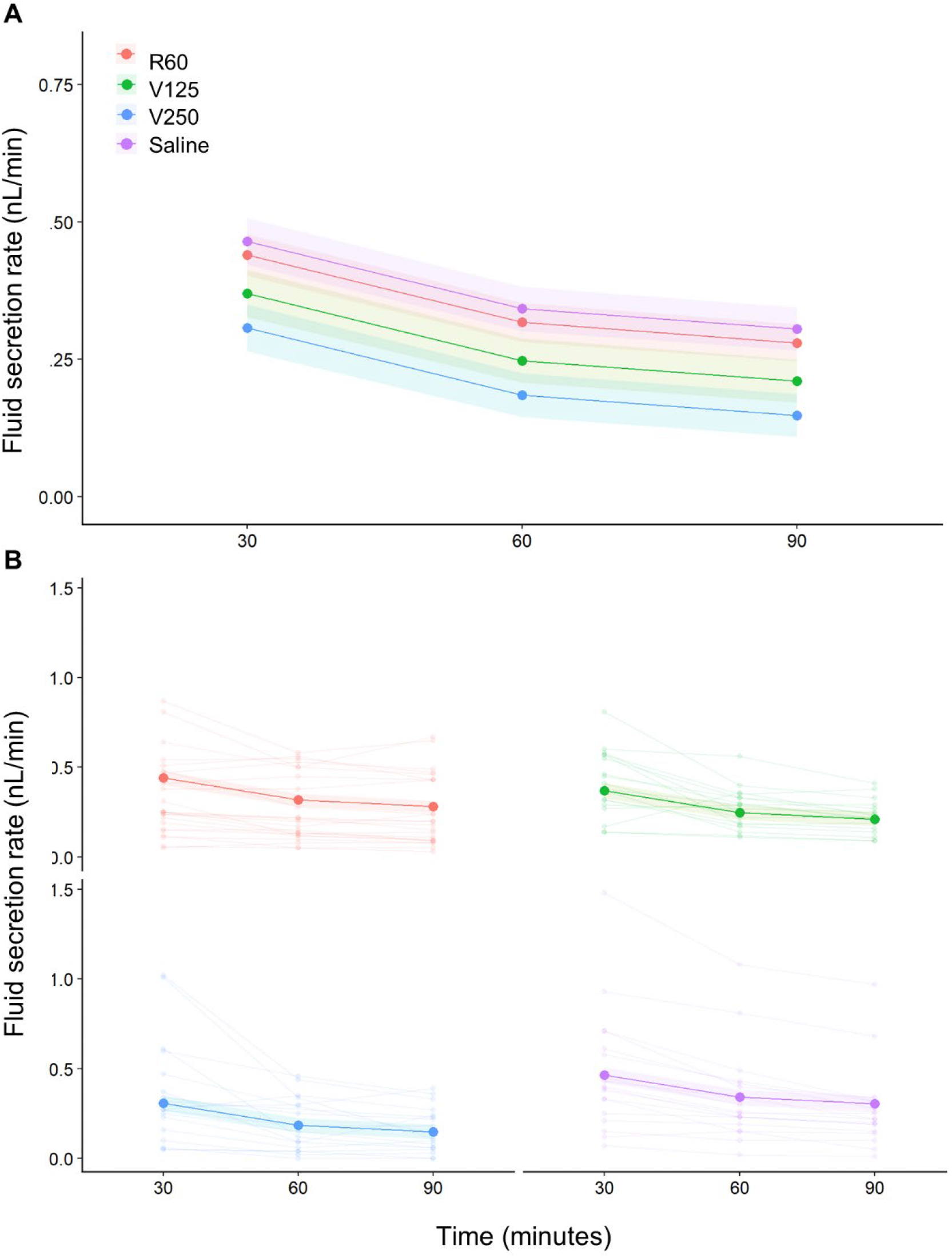
Mean fluid secretion rate of Malpighian tubules during the incubation period. (**A**) The mean fluid secretion rate decreased after 60 minutes of incubation in all the treatments, while it remained steady for the next 30 minutes in all the treatments except in the saline one. Each line with the shaded area around represents the mixed effect model fit with the standard error. (**B**) As in ‘A’, but with the fluid secretion rate of individual tubules shown by pale lines linking smaller dots.

### Renewing bath saline increases the fluid secretion rate

To exclude the possibility that the decrease in the fluid secretion rate was caused by a damage of the tubules during the Ramsay assay, we replaced the saline bath after 90 minutes with fresh saline, to determine whether tubules would increase their fluid secretion rate to previous levels. We removed three tubules from six locusts (17 tubules in total, one tubule excluded) and incubated them in saline for 90 minutes, removing the droplet secreted every 30 minutes. After 90 minutes the saline bath was removed, replaced with fresh saline. The tubules were then incubated for further 90 minutes removing the droplet secreted every 30 minutes. The secretion rate decreased after 60 and 90 minutes (S4 Fig, S2 Table), but increased after 120 minutes following replacement of the saline (S4 Fig, S2 Table).

### Malpighian tubule integrity during the Ramsay assay

To exclude the possibility that manipulation during the Ramsay assay altered the diameter of the tubules, we measured the diameter of the tubules *in vivo*, at the beginning, and at the end of the assay. We found that the diameter of the tubules was unaffected by the assay and was comparable to the tubule’s diameter *in vivo* (S5 Fig, S3A,B Table).

### Net extrusion of Rhodamine B

We determined the concentration of Rhodamine B in each of the droplets collected from the Malpighian tubules exposed to the R60, V125 and V250 treatments at each time point. There was a significant interaction between treatment and time (F_4,73.19_=15.19, p<.001, Fig 4A), indicating that the rhodamine B concentration changed over time depending on the treatment (Table 2). The concentration of rhodamine B in the droplets significantly increased during the incubation time in all the treatments (Table 1B, Fig 4A). In particular, the increase in rhodamine B concentration in the R60 treatment was more pronounced than in either the V125 or V250 treatments (Table 1C, Fig 4A). At each time point, the treatment significantly affected the rhodamine B concentration. Compared to the R60 treatment, the addition of verapamil in the V125 and V250 treatments reduced the rhodamine B concentration of the secreted droplets after 30, 60, and 90 minutes (Table 2D, Fig 4A). However, there was no significant difference in the rhodamine B concentration between the V125 and V250 treatments at any time point (Table 2D, Fig 4A).

**Fig 4.**
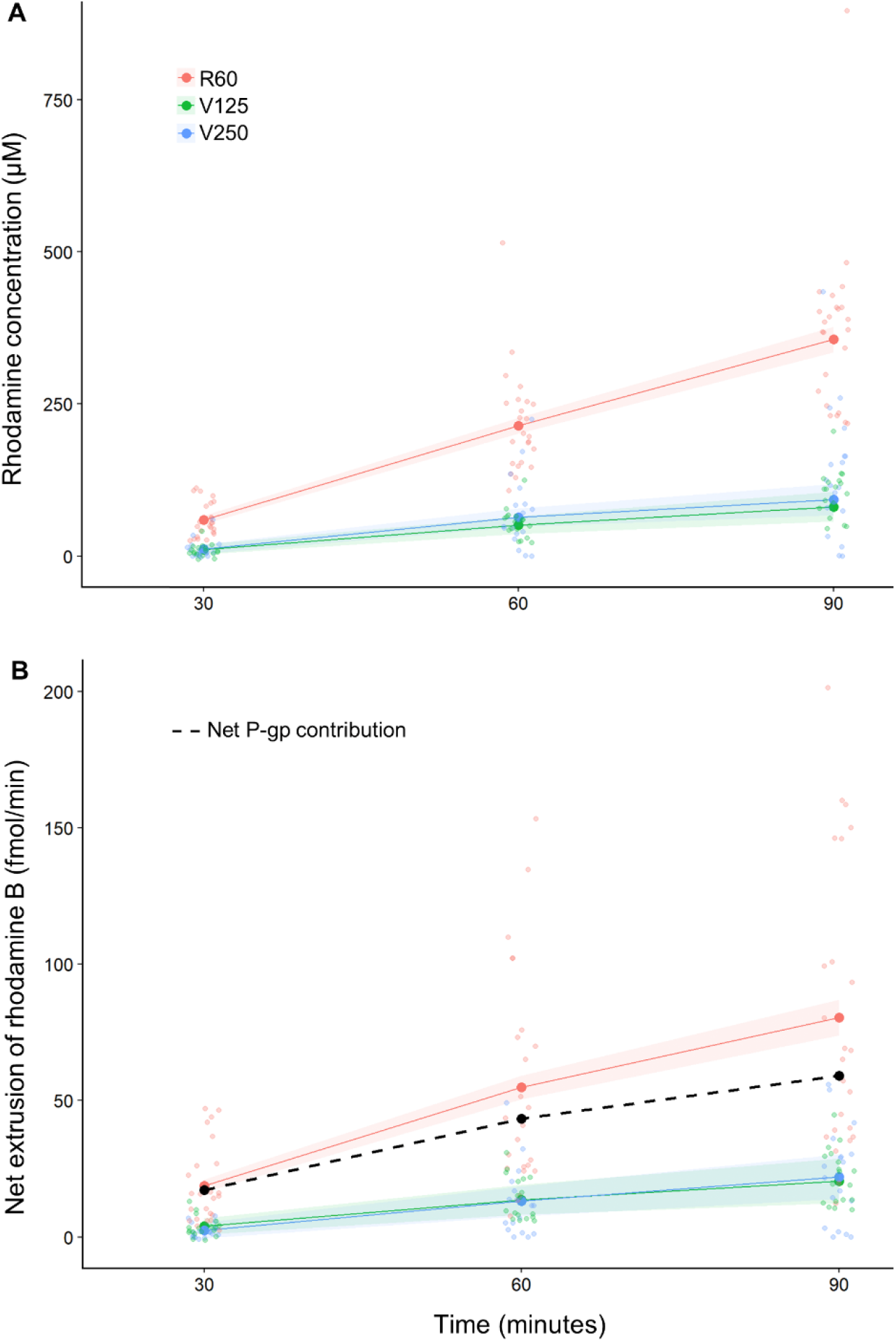
The mean rhodamine B concentration of the droplets secreted by the Malpighian tubules, and the mean net extrusion of rhodamine B increased over time in all the treatments. (**A**) The addition of verapamil (125 or 250 μM) reduced the rhodamine B concentration of the droplets secreted compared with the control treatment 60 μM rhodamine. (**B**) The addition of verapamil (125 μM or 250 μM) reduced the net extrusion of rhodamine B compared with the control treatment 60 μM rhodamine. The dashed line represents the net contribution of the P-glycoproteins obtained by subtracting the net extrusion of rhodamine in the V250 treatment from the R60. Each line with shaded region represents the mixed effect model fit with the standard error. Small circles represent the raw data.

**Table 2.** The outcomes of linear mixed effect model used to investigate the effect of incubation time and treatment on the rhodamine concentration of the droplets secreted.

| | | Treatment | Time 30<br>Mean ± se | Time 60<br>Mean ± se | Time 90<br>Mean ± se |
| --- | --- | --- | --- | --- | --- |
| A | Rho concentration (μM) | R60 | 59.0 ± 7.3 | 217.9 ± 13.7 | 369.7 ± 22.4 |
| | | V125 | 9.0 ± 8.5 | 54.0 ± 15.8 | 103.4 ± 25.9 |
| | | V250 | 10.4 ± 8.5 | 74.4 ± 15.9 | 135.2 ± 25.9 |

|  | Treatment | Time 60 vs time 30 |  | Time 90 vs time 60 |  |
| --- | --- | --- | --- | --- | --- |
| | | estimate ± se | P-value | estimate ± se | P-value |
| B | R60 | 158.9 ± 11.8 | <.001 | 151.8 ± 11.8 | <.001 |
| | V125 | 45 ± 13.6 | 0.004 | 49.4 ± 13.6 | 0.001 |
| | V250 | 64 ± 13.7 | <.001 | 60.8 ± 13.7 | <.001 |
| C | V125 vs R60 | -113.9 ± 18 | <.001 | -102.4 ± 18 | <.001 |
| | V250 vs R60 | -94.9 ± 18.1 | <.001 | -91 ± 18.1 | <.001 |
| | V250 vs V125 | 19 ± 19.3 | 0.588 | 11.4 ± 19.3 | 0.826 |

|  | Treatment | Time 30 |  | Time 60 |  | Time 90 |  |
| --- | --- | --- | --- | --- | --- | --- | --- |
| | | estimate ± se | P-value | estimate ± se | P-value | estimate ± se | P-value |
| D | V125 vs R60 | -49.98 ± 11.2 | <.001 | -163.88 ± 20.9 | <.001 | -266.31 ± 34.2 | <.001 |
| | V250 vs R60 | -48.59 ± 11.2 | <.001 | -143.49 ± 21.0 | <.001 | -234.53 ± 34.2 | <.001 |
| | V250 vs V125 | 1.39 ± 12.0 | 0.993 | 20.4 ± 22.5 | 0.637 | 31.78 ± 36.6 | 0.662 |

The model applied was (Rhodamine concentration ∼ time \* treatment + (1|locust) + (1+time|tubule)).
(A) Summary of the mean values for each treatment at each time point. (B) Pairwise comparisons between time of incubation separately for each treatment. (C) Pairwise comparison of the interaction between time and treatment. D) Pairwise comparisons between treatments separately for each time of incubation.

We calculated the total number of moles of rhodamine B extruded by the Malpighian tubules per minute as the product of the secretion rate and the rhodamine B concentration of the droplets secreted, which we termed the net extrusion rate. There was an interaction between treatment, time and surface (F_4,74.59_=2.56, p=0.046; Table 3, Figs 4B,5A-C), indicating that the dependency of the net extrusion of rhodamine B upon tubule surface area was influenced by the incubation time and the treatment (Table 3A,E, Fig 5A-C). The net extrusion increased significantly between 30 and 60 minutes in all the treatments (Table 3B, Fig 4B). In contrast, between 60 and 90 minutes the net extrusion increased only for the tubules incubated in R60 and V250, whereas it remained steady for V125 treatments (Table 3B, Fig 4B). In particular, the net extrusion was more pronounced in the R60 treatment compared to V125 and V250 (Table 3C, Fig 4B). At each time point, the treatment affected the net extrusion of rhodamine B (Table 3D, Fig 4B). In comparison to the R60 treatment, the addition of verapamil in the V125 and V250 treatments significantly reduced the net extrusion of rhodamine B after 30, 60 and 90 minutes, however, there was no significant difference between the V125 and V250 treatments (Table 3D, Fig 4B).

**Table 3.**
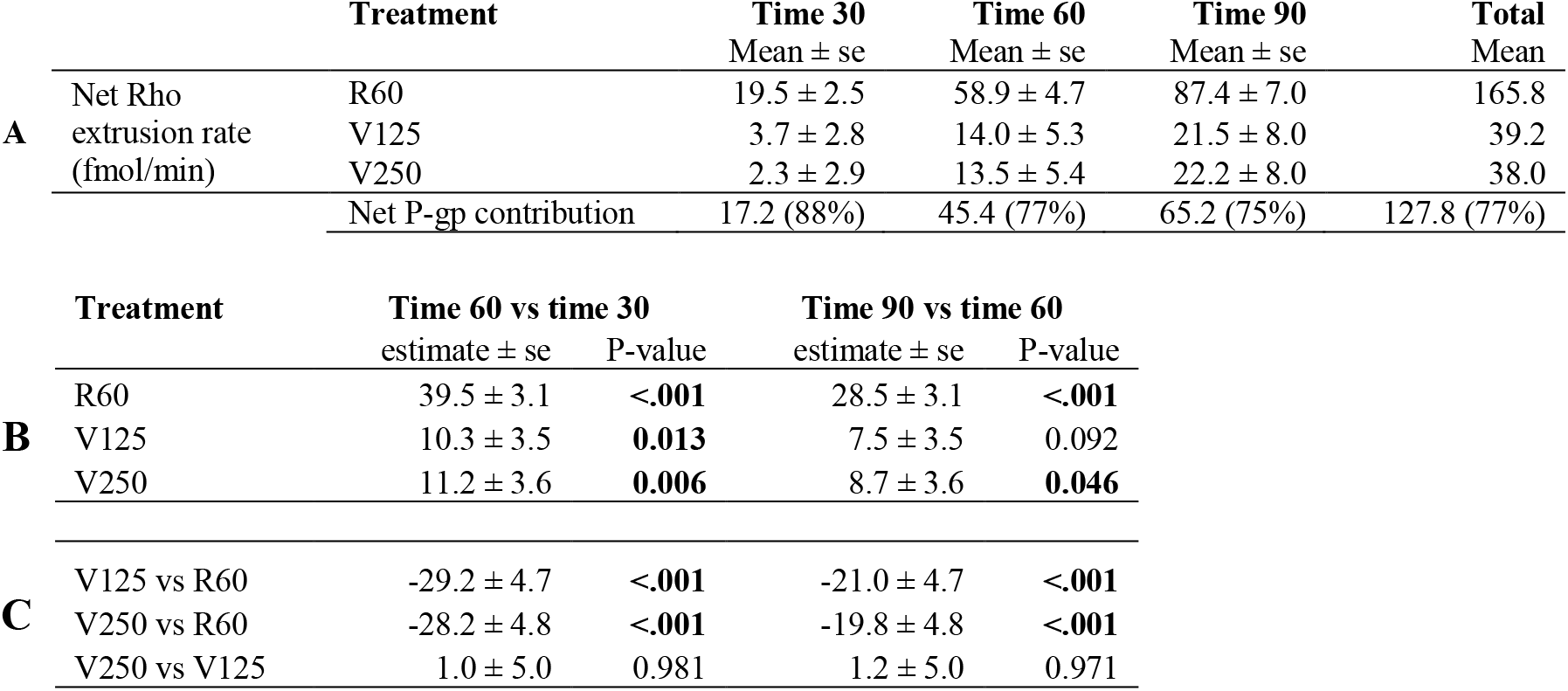
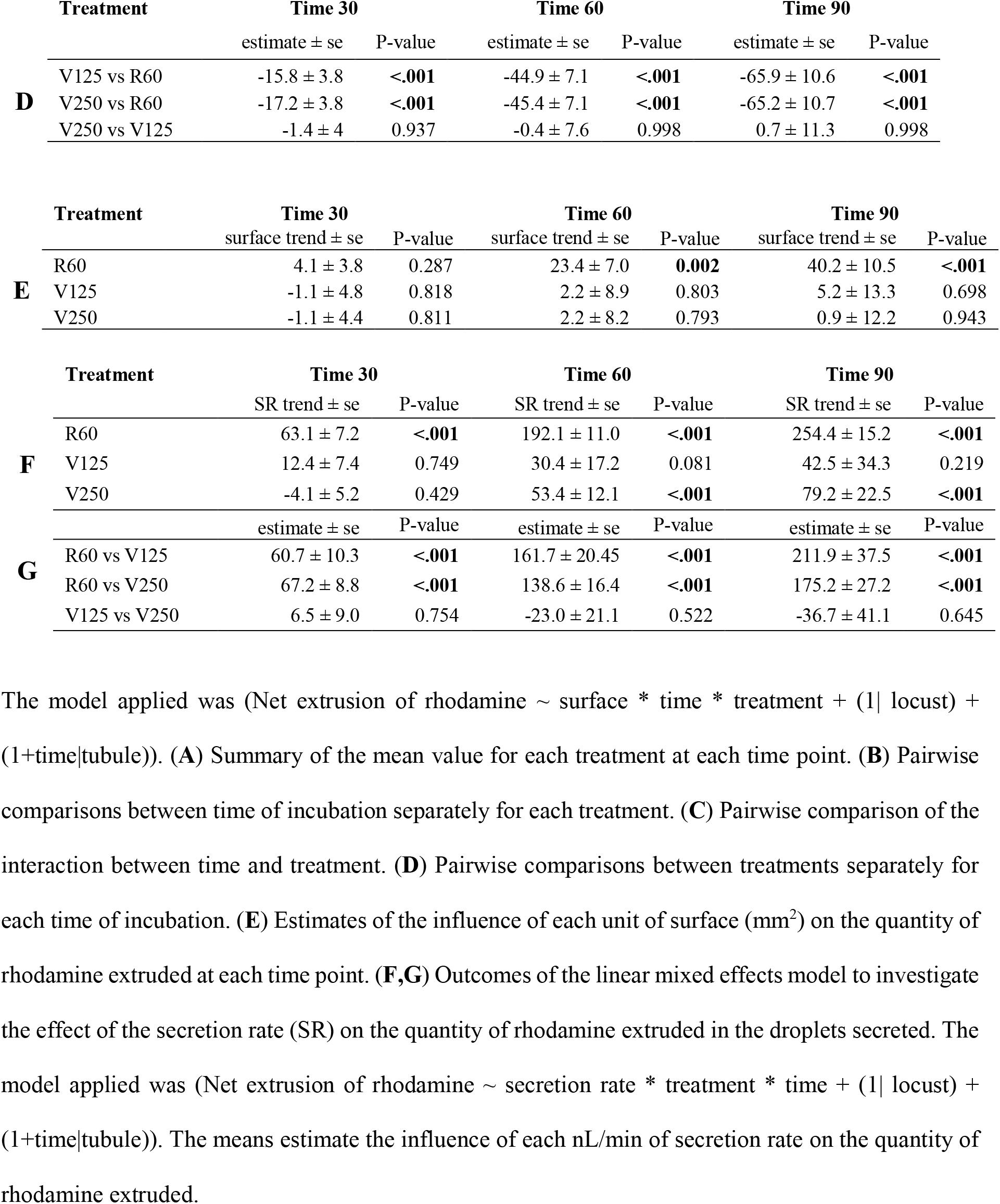
Outcomes of a linear mixed model to investigate the effect of incubation time and treatment on the net quantity of rhodamine extruded in the droplets secreted per minute.

**Fig 5.**
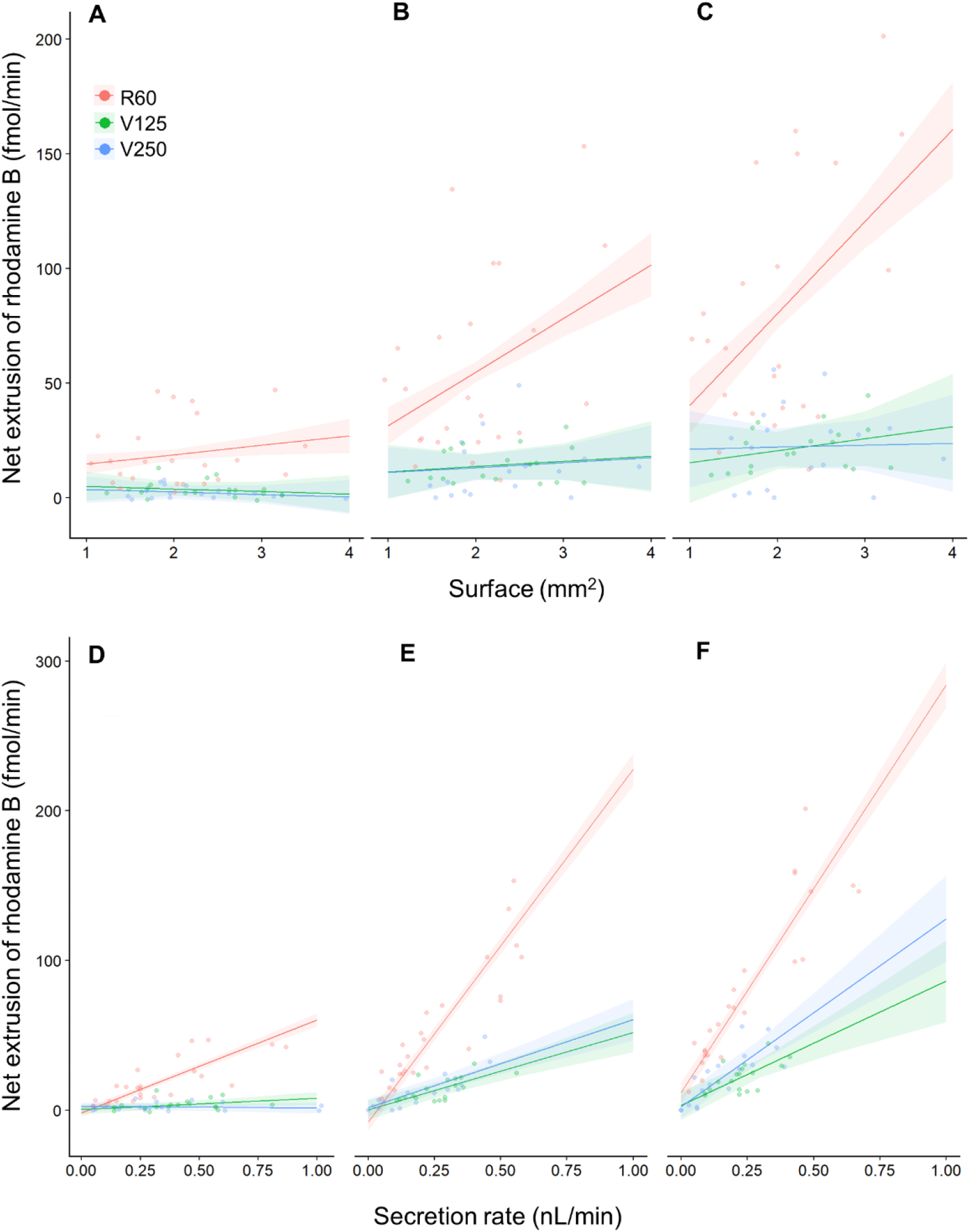
Tubule surface area and the fluid secretion rate positively influences the net extrusion of rhodamine B. (**A**) A plot of the surface area of the tubule exposed to the bath versus the net extrusion of rhodamine B for each of the three treatments after 30 minutes of incubation. (**B**) As in ‘A’ but after 60 minutes. (**C**) As in ‘A’ but after 90 minutes. (**D**) A plot of the fluid secretion rate versus the net extrusion of rhodamine B for each of the three treatments after 30 minutes of incubation. (**E**) As in ‘D’ but after 60 minutes. (**F**) As in ‘D’ but after 90 minutes. Small circles indicate the net extrusion of rhodamine B of individual tubules at a particular time point. Each line with the shaded area around represents the mixed effect model fit with the standard error.

The surface area positively influenced the net extrusion of rhodamine B in the tubules incubated for 60 and 90 minutes in the R60 treatment, while there was no correlation between surface area and net extrusion in V125 and V250 treatments (Table 3E, Fig 5A-C). In addition, we found that after 30 minutes the fluid secretion rate of the tubules incubated in R60 positively influenced the net extrusion of rhodamine B, while there was no significant effect in the V125 and V250 treatments (Table 3F, Fig 5D). After 60 and 90 minutes, the fluid secretion rate positively correlated with the net extrusion of rhodamine B in R60 and V250 treatments, but not V125 (Table 3F, Fig 5E,F). Moreover, the secretion rate of the tubules incubated in R60 showed a more pronounced effect on the net extrusion of rhodamine B than that of the tubules incubated in V125 and V250 (Table 3G, Fig 5D-F).

### Net contribution of P-glycoproteins

A single Malpighian tubule with a mean surface area incubated in a solution of 60 μM rhodamine B, extruded rhodamine B with a rate of 19.5 ± 2.6 fmol/min in the first 30 minutes, 58.9 ± 4.7 fmol/min between 30 and 60 minutes and 87.4 ± 7.0 fmol/min between 60 and 90 minutes (Table 3A). The addition of verapamil significantly reduced the quantity of rhodamine B extruded but did not block it completely (Table 3A). The rhodamine B concentration of the droplets secreted did not differ between the V125 and V250 treatments (Table 2A). Therefore, we assumed that 125 μM verapamil was sufficient to inhibit all the P-glycoproteins that are verapamil sensitive and that passive diffusion and other types of pumps may contribute to extrusion of Rhodamine B. We subtracted the mean values of the net extrusion of rhodamine B obtained from the V250 treatment from the net extrusion of the R60 treatment to estimate the contribution of the P-glycoproteins alone. The average rates of rhodamine B extruded by the P-glycoproteins were 17.2 fmol/min, 45.4 fmol/min and 65.2 fmol/min between 0-30, 30-60 and 60-90 minutes respectively (Table 3A, Fig 4B dashed line). In percentage, the P-glycoproteins are responsible for the 88%, 77% and 75% (between 0-30, 30-60 and 60-90 minutes respectively) of the total extrusion of rhodamine B. Overall, P-glycoproteins account for the 77% of the total extrusion of rhodamine B, over the 90 minutes of incubation.

## Discussion

Our aim was to determine whether P-glycoprotein transporters are involved in the removal of xenobiotic substances by Malpighian tubules from the haemolymph of desert locusts and, if so, how they perform physiologically. To this end, we developed an alternative method to liquid chromatography–mass spectrometry [37], radiolabelled alkaloids [9] or confocal microscopy [11,12] based upon measuring dye concentration. A similar method has been used previously to investigate epithelial transport in tardigrades and desert locusts using chlorophenol red by imaging through the gut or tubules [38]. By imaging extruded drops from Malpighian tubules, we assessed the performance of epithelial transporters more accurately than by imaging the lumen of the tubules. We used the P-glycoprotein substrate rhodamine B [25] and the P-glycoprotein inhibitor verapamil [39] to assay P-glycoprotein function through the colour of the droplets secreted by the Malpighian tubules. Using this strategy, we obtained evidence that desert locust Malpighian tubules express a P-glycoprotein transporter, that the fluid secretion rate of these tubules is proportional to their surface area, and that the fluid secretion rate influences the net extrusion of rhodamine B.

### A P-glycoprotein transporter extrudes xenobiotics in desert locust Malpighian tubules

Our conclusion that desert locust Malpighian tubules express P-glycoproteins is supported by two lines of evidence. Firstly, these tubules actively extrude the dye rhodamine B, a P-glycoprotein substrate (e.g. [25,26]), when it is present in the solution in which they are incubated. Rhodamine B has been widely used as a substrate for P-glycoproteins in cell culture and blood brain barrier models (e.g. [25,26]). Secondly, the addition of verapamil, a P-glycoprotein inhibitor (e.g. [23,27,28,39]), significantly reduced the extrusion of rhodamine B. The presence of P-glycoproteins in other tissues of the desert locusts also supports our conclusion that they are expressed in the Malpighian tubules; proteins with a comparable physiology and similar sequence to the human P-glycoprotein (ABCB1 gene) are expressed within the blood brain barrier of the locusts *S. gregaria* and *L. migratoria* [23,37,40,41]. Further support comes from comparison with other insects; P-glycoprotein transporters have been found in the Malpighian tubules of many other insects including the black field cricket (*Teleogryllus commodus* [11]), tobacco hornworm (*Manduca sexta* [9,42]), fruit fly (*D. melanogaster*), kissing bug (*Rhodnius prolixus*), large milkweed bug (*Oncopeltus fasciatus*), yellow fever mosquito (*Aedes aegypti*), house cricket (*Acheta domesticus*), migratory locust (*Locusta migratoria*), mealworm beetle (*Tenebrio molitor*), and the American cockroach (*Periplaneta americana*) [15].

Rhodamine B extrusion was not blocked entirely by 125 μM verapamil or by double this concentration. Comparison of the extent of the reduction at both concentrations suggests that even at the lower verapamil concentration all the P-glycoproteins were inhibited. This suggests that approximately 77% of rhodamine B was transported by P-glycoproteins via a verapamil sensitive mechanism, whilst the remaining 23% was verapamil insensitive. Moreover, after 90 minutes the concentration of rhodamine B in the droplets secreted by the Malpighian tubules was higher than that of the bathing solution suggesting that rhodamine B transport in the presence of verapamil is not simply due to passive diffusion, but that other active transporters may be implicated. Several potential candidates for alternative active transporters exist. For example, in human cell lines (Calu-3), rhodamine B can interact with multiple organic cation transporters (OCT3, OCTN1,2) [43]. Potentially, however, verapamil may be incapable of blocking rhodamine B transport completely allowing a small number of rhodamine B molecules to be extruded by the P-glycoprotein even in the presence of verapamil.

Verapamil inhibits P-glycoproteins and does not interact with other multidrug resistance proteins [44]. This suggests that the effects of verapamil in our experiments are through its specific effects upon P-glycoproteins. The mechanistic basis of verapamil inhibition is unclear but the most widely accepted explanation is that P-glycoproteins extrude both verapamil and their substrate but that verapamil diffuses back across the lipid bilayer much faster than the substrate creating a futile cycle and thereby competing with the substrate transport [45,46].

Verapamil is, however, also a known L-type Ca^2+^ channel blocker [47]. In *Drosophila*, L-type Ca^2+^ channels are expressed in the basolateral and apical membranes of the tubule principal cells, and are involved in the regulation of the fluid secretion [48]. Indeed, an increase of the intracellular Ca^2+^ level mediates the effect of diuretic hormones [49]. Verapamil reduced the fluid secretion of *Drosophila* Malpighian tubules stimulated by peptide agonists (e.g. CAP2b, cGMP) but had no effect on unstimulated tubules [48]. Our experiments cannot exclude the possibility that verapamil affects Ca^2+^ channels in locust Malpighian tubules by interfering with intracellular Ca^2+^, and thereby has an indirect effect on Rhodamine B extrusion. However, because the fluid secretion rate did not differ between treatments, the reduction in the net extrusion of rhodamine B was likely caused solely by verapamil inhibition of the P-glycoproteins. Thus, when considered within the context of the expression of P-glycoproteins in the blood brain barrier of locusts [23,37,40,41] and the expression of P-glycoproteins in homologous Malpighian tubules in other insects species (see above), it seems highly likely that the tubules of gregarious desert locust express P-glycoproteins.

### Malpighian tubule surface area and fluid secretion rate

We used linear mixed effect models to determine which physical feature of a tubule, its surface area, length or diameter has the greatest influence on fluid secretion rate. We found that the surface area of the Malpighian tubules positively influences their fluid secretion rate, and more accurately predicts the fluid secretion rate than tubule diameter or length. Some previous studies have reported that tubule length was linearly related to the fluid secretion rate [50,51] but in our analysis length was consistently worse than surface area as a predictor. Even when the surface area (or length) has been accounted for in previous studies, this has involved dividing the fluid secretion rate by the surface area (or length) to obtain the fluid secretion rate per unit area (or unit length) [15,52]. Although this approach is useful when comparing tubules of different sizes from different insect species, it fails to reveal the exact relationship between the surface area and the fluid secretion rate. Our statistical models demonstrate that interactions between factors such as surface area and fluid secretion rate depend upon the treatment applied. For example, our results show that the fluid secretion rate increases with the increasing of the tubule surface in all the treatments apart from 250 μM verapamil, where the surface no longer influences the fluid secretion rate. Such interdependencies are unlikely to be detected or accounted for in simpler statistical analyses, causing such interdependencies to be ignored.

### Reliability of isolated Malpighian tubule measurements

The decrease in fluid secretion rate that we found is comparable with other studies of isolated Malpighian tubules, where the fluid secretion rate of tubules incubated in saline decreased of 30% after 20 minutes [53] or over one hour [54]. In contrast, fluid secretion rates have been found steady over time in studies where the Malpighian tubules were left attached to the gut through the tracheae, and the whole preparation immersed in saline [6,55]. One of the differences between the isolated tubule assay and the whole gut assay is the shorter portion of trachea in contact with the tubule immersed in the bathing solution. Therefore, the reduction of the fluid secretion rate in isolated tubules may be a consequence of a smaller amount of residual oxygen in the tracheae compared with the whole gut preparation. Additionally, the volume of the bathing solution in the isolated tubule assay is far smaller than in the whole gut preparation, which could lead to a quicker depletion of oxygen and/or other substances in the bathing solution. Replacing the bathing solution with fresh saline produced an increase in the fluid secretion rate, supporting this interpretation. Finally, there could be a decrease in the tubule diameter during the assay compared to the *in vivo* preparation. We can safely exclude this, however, because we found that the tubule diameter was unaffected by the assay and was similar to the tubule’s diameter *in vivo*.

### Determining the transepithelial transport of xenobiotics by P-glycoprotein transporters in locust Malpighian tubules

Transepithelial transport of xenobiotics by P-glycoproteins across the inner membrane of the Malpighian tubules can be determined from the net extrusion rate we measured. The disparity between the transepithelial transport and the net extrusion rate is due to the properties of rhodamine B. This lipophilic dye can passively permeate the lipid bilayer of liposomes, following the concentration gradient [25]. Likewise, in desert locust Malpighian tubules, rhodamine B can back diffuse into the bath solution when active transporters increase its luminal concentration creating a concentration gradient. Consequently, the fluid secretion rate affects the net extrusion rate of rhodamine B because a decrease in the fluid secretion will increase the luminal concentration of the dye, allowing greater back diffusion whilst an increase in the fluid secretion rate will have the opposite effect [1]. Therefore, if a dye, like rhodamine B, can back diffuse, its net extrusion will be proportional to the fluid secretion rate (S6A-C Fig), while if the dye cannot back diffuse there will be no correlation (S6D-F Fig). When the rhodamine B concentration in the lumen was lower or similar to the bath (60 μM) the net transport would be little affected by the secretion rate. Conversely, when the luminal rhodamine B concentration was higher than the bathing solution, part of the dye diffused back, so that the net extrusion was significantly influenced by the fluid secretion rate.

Previous studies demonstrated that Malpighian tubules can exhibit different degrees of passive permeability to different substances, depending on the species and on the properties (i.e. size, polarity) of the substrate used (e.g. dyes, alkaloid, drugs) [1]. For example, tubules of the kissing bug (*R. prolixus)* have passive permeability to the alkaloid nicotine [8], but not to the dye indigo carmine [1], whereas tubules of the blowfly (*Calliphora erythrocephala)* are permeable to indigo carmine [1]. Thus, when studying the effect of a substance on active transporters, it is important to take into account the fluid secretion rate because a change in the net extrusion rate of a substrate may be caused not only by a direct effect on the transporters, but also by an indirect consequence of a change in the fluid flow [12]. In our experiment, the fluid secretion rate did not differ between treatments, indicating that the reduction of net extrusion of rhodamine B following exposure to verapamil was caused solely by inhibition of the P-glycoprotein. However, the fluid secretion rate decreased over time, which may produce an underestimation of the net transepithelial transport of rhodamine B.

### Implications for desert locust detoxification

Gregarious desert locusts feed on a broad variety of plants including species containing secondary metabolites such as atropine to become unpalatable to predators [17–20]. The expression of P-glycoproteins in the Malpighian tubules to extrude noxious substances may be an adaptation to cope with the ingestion of toxic plants. This may also be the reason for expression of P-glycoproteins on the blood brain barrier of desert locusts, which would prevent the uptake of hydrophobic substances in the central nervous system [23]. Yet the relationship between ingesting toxins and detoxification pathways in the Malpighian tubules is not straightforward; some species of Orthoptera, as well as Coleoptera, Lepidoptera, Heteroptera, Hymenoptera and Sternorrhyncha [56], sequester toxins from the plants they ingest to deter predators. However, toxicity may also be conferred by gut contents, rather than through sequestration within bodily tissues. For example, the chemical defence of the spotted bird grasshopper, *Schistocerca emarginata* (*=lineata*), is mediated by the contents of toxic plant in its gut [21,22]. This species is a congener of the desert locust, *S. gregaria*, which suggests that a similar strategy may be involved in the production of toxicity in this species. If this is the case, then the presence of toxins in the haemolymph may be a consequence of ingesting toxic plant material for storage within the gut. In such a scenario, detoxification pathways within the Malpighian tubules would then be essential for ensuring that toxins do not accumulate within the haemolymph to concentrations that would affect physiological processes.

The P-glycoprotein detoxification pathway that we have characterised in desert locusts is likely to be highly effective in extruding xenobiotic compounds from haemolymph, especially when the number of Malpighian tubules within an individual locust is considered. However, it is important to consider that the locusts used for our experiments have experienced a diet free of toxins, such as atropine. P-glycoprotein expression can be modulated depending on the diet [57]; *Drosophila* larvae fed on colchicine increased the expression of the P-glycoprotein gene homologue mdr49 in the brain and gut. Consequently, adult gregarious desert locusts that have fed on a diet including plant toxins may have even stronger P-glycoprotein detoxification pathways. In contrast to their gregarious counterparts, solitary desert locusts actively avoid plants containing atropine [17], and find odours associated with it aversive [58]. Thus, Malpighian tubules of solitary desert locusts may express fewer P-glycoproteins than those of the gregarious phase, possibly due to their reduced exposure to secondary metabolites from their diet.

### Implications for insect pest control

P-glycoproteins are implicated in the resistance of some insect pests to insecticides [59] promoting the efflux of various xenobiotics thereby decreasing the intracellular drug accumulation. For instance, P-glycoproteins have been detected in a resistant strain of the pest cotton bollworm (*Helicoverpa armigera*) but not in a susceptible one [60]. P-glycoproteins have also been implicated in the protection of the mitochondria from insecticide damage [61]. In Africa and Asia, applications of insecticides are carried out to try to control desert locust plagues [62]. To increase the efficacy, a combination of P-glycoprotein inhibitors and insecticide may act synergistically to increase the locust mortality, reducing the amount of insecticide used. Further investigation into the interaction between P-glycoproteins and different xenobiotics (i.e. insecticides, herbicides, miticides and secondary metabolites) may improve our understanding of the physiological effect of pesticides on insects, and subsequently lead to the development of more specific targeted insecticides.

## Acknowledgements

M.R. was financially supported by a graduate studentship from the School of Life Sciences, University of Sussex. D.D.B was financially supported by the Welsh government/HEFCW RESILCOAST project.

## Author contributions

M.R. and J.E.N conceived the experiment; M.R. and J.E.N designed the experiments. M.R. carried out the experiments, analysed the images and carried out the data analyses. D.D.B provided help for the statistical analyses. M.R. and J.E.N wrote the manuscript.

## Supporting information

**S1 Fig.**
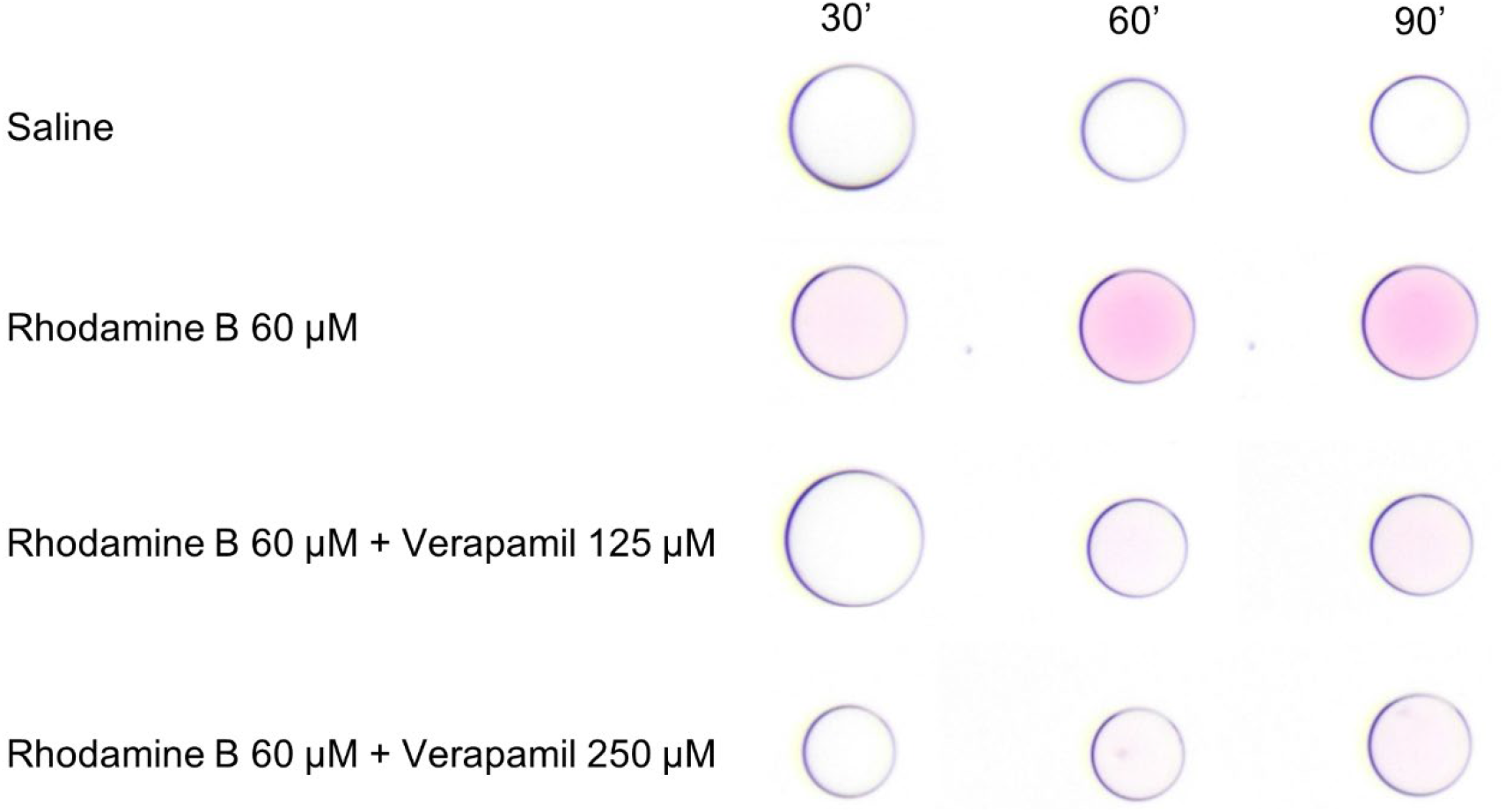
Examples of the droplets secreted by the Malpighian tubules of desert locusts during the incubation in each of the different treatments every 30 minutes. The size of each droplet depends upon the fluid secretion rate whilst the colour is determined by the net extrusion rate of rhodamine B.

**S2 Fig.**
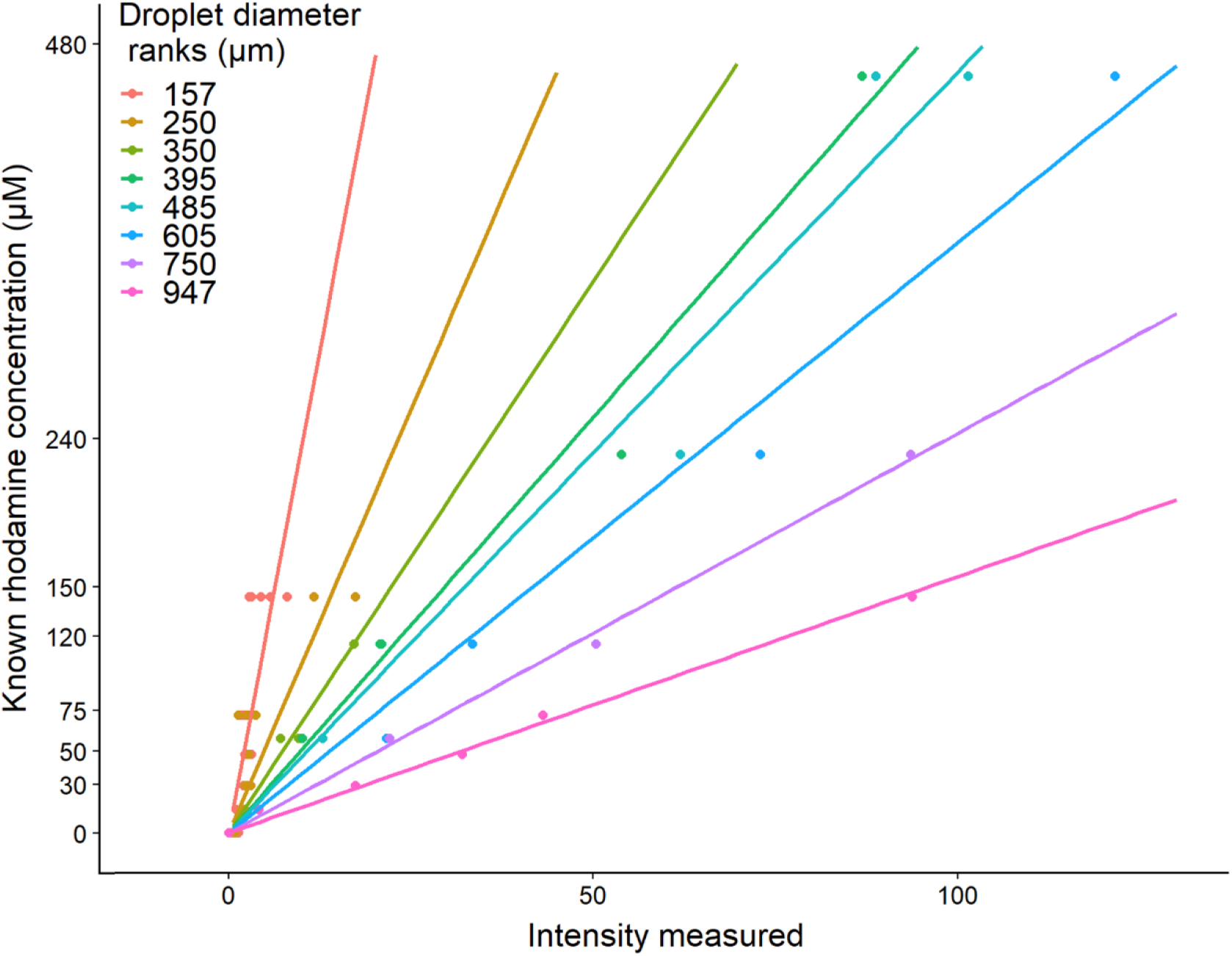
Calibration lines used to determine the rhodamine B concentration in the droplets secreted by the Malpighian tubules. For each droplet diameter rank there is a linear relationship between the rhodamine concentration of the droplet and the colour intensity measured. The slope of the lines decreases as the diameter increases. We estimated the rhodamine B concentration of the droplets secreted by measuring the colour intensity and the diameter of each droplet. Each line represents the linear regression fit for each mean diameter rank.

**S3 Fig.**
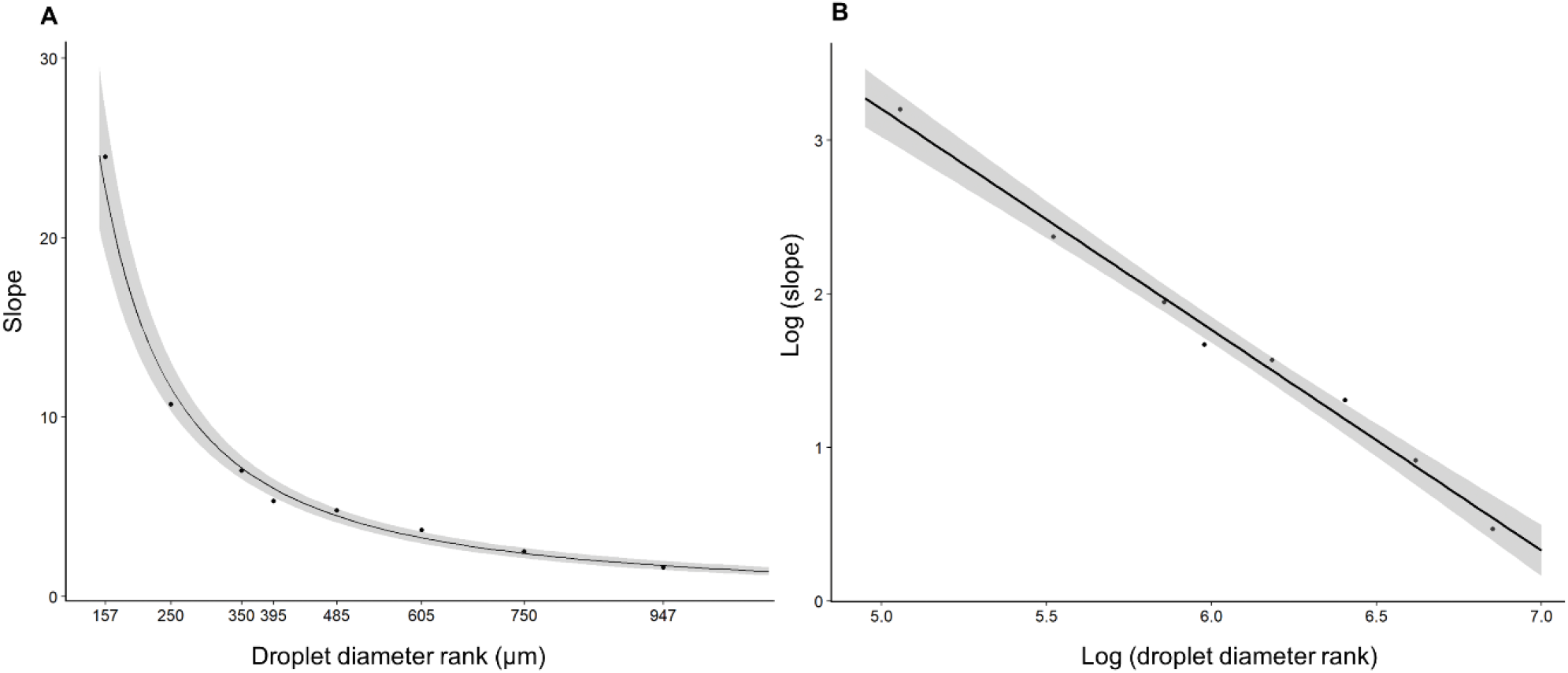
The relationship between intensity and rhodamine B concentration depends upon droplet diameter. (A) The slope decreases as the diameter increases, following an exponential decay. (B) After log transformation the relationship becomes linear. Using this linear equation for each droplet diameter measured, we predicted the slope of the line equation that link the colour intensity to the rhodamine concentration.

**S4 Fig.**
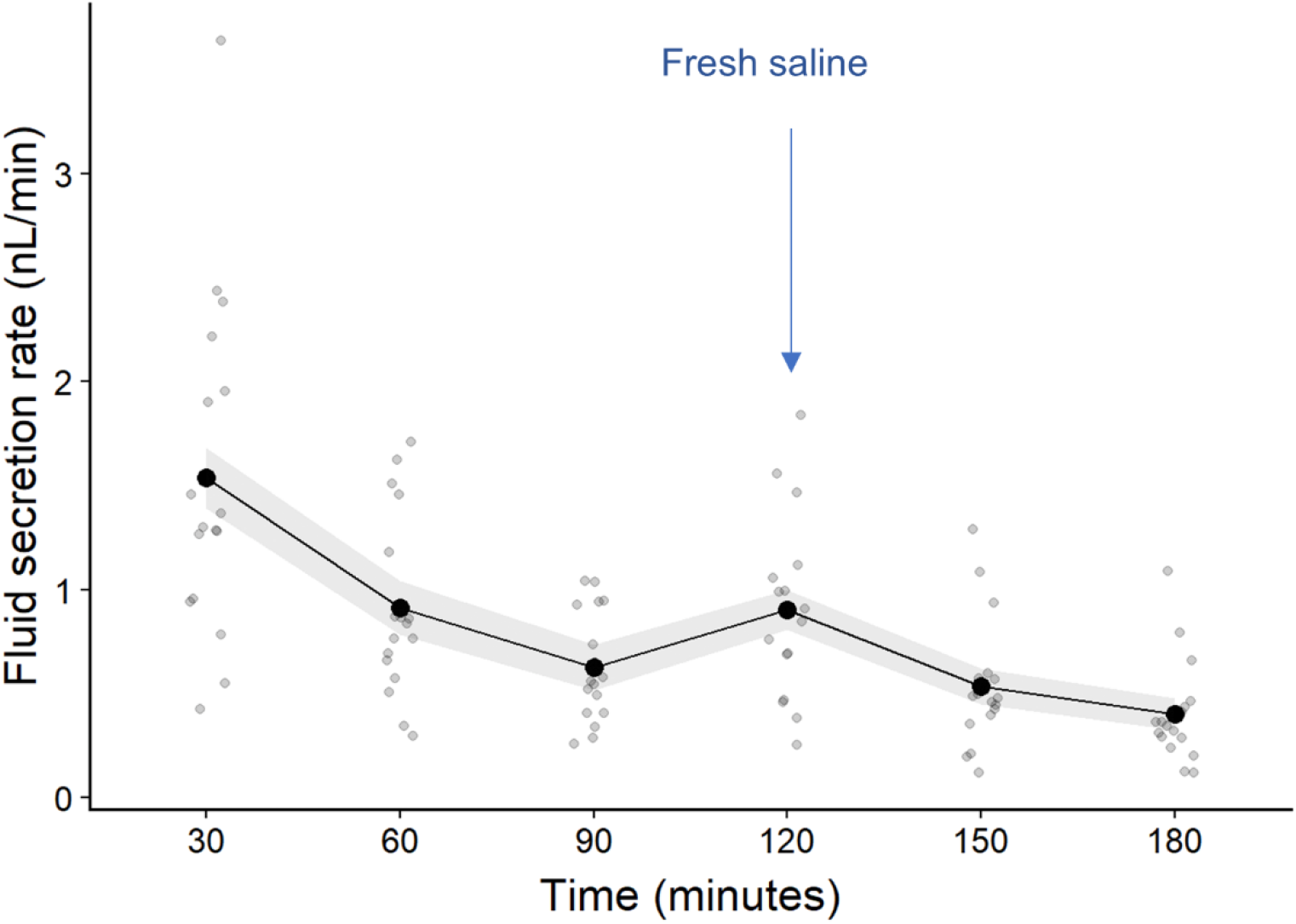
Mean fluid secretion rate of Malpighian tubules increases with fresh saline. The tubules were incubated in saline but after 90 minutes the saline bath was removed and replaced with fresh saline. The arrow indicates the first measurement taken after the saline had been replaced. Grey points indicate the fluid secretion rate of individual tubules at a particular time point.

**S5 Fig.**
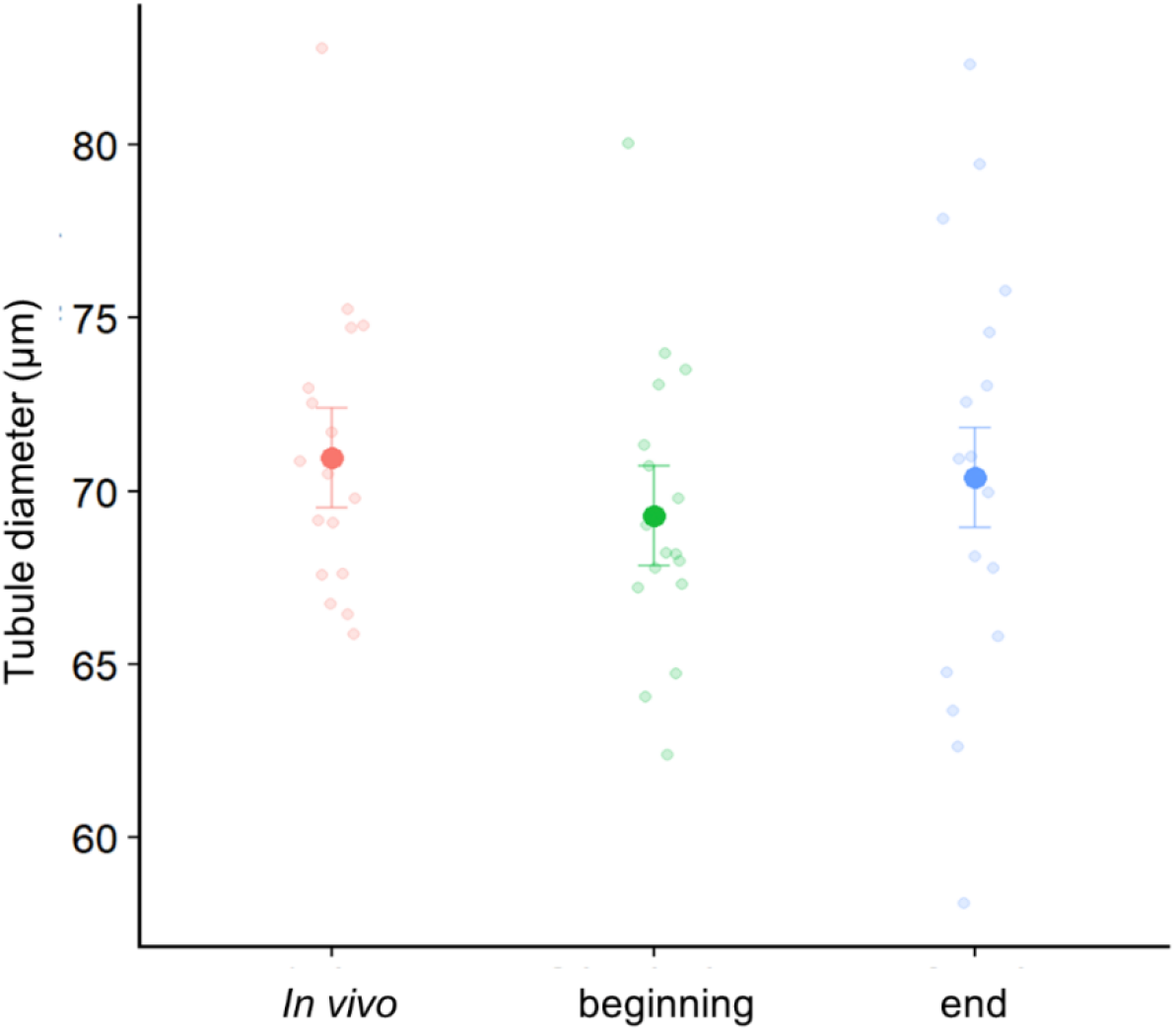
Manipulation of Malpighian tubules in the Ramsay assay did not affect their diameter. To exclude the possibility that manipulation during the assay affected tubule morphology, we measured the tubule’s diameter *in vivo*, at the beginning, and at the end of the assay. The diameter was unaffected by the manipulation.

**S6 Fig.**
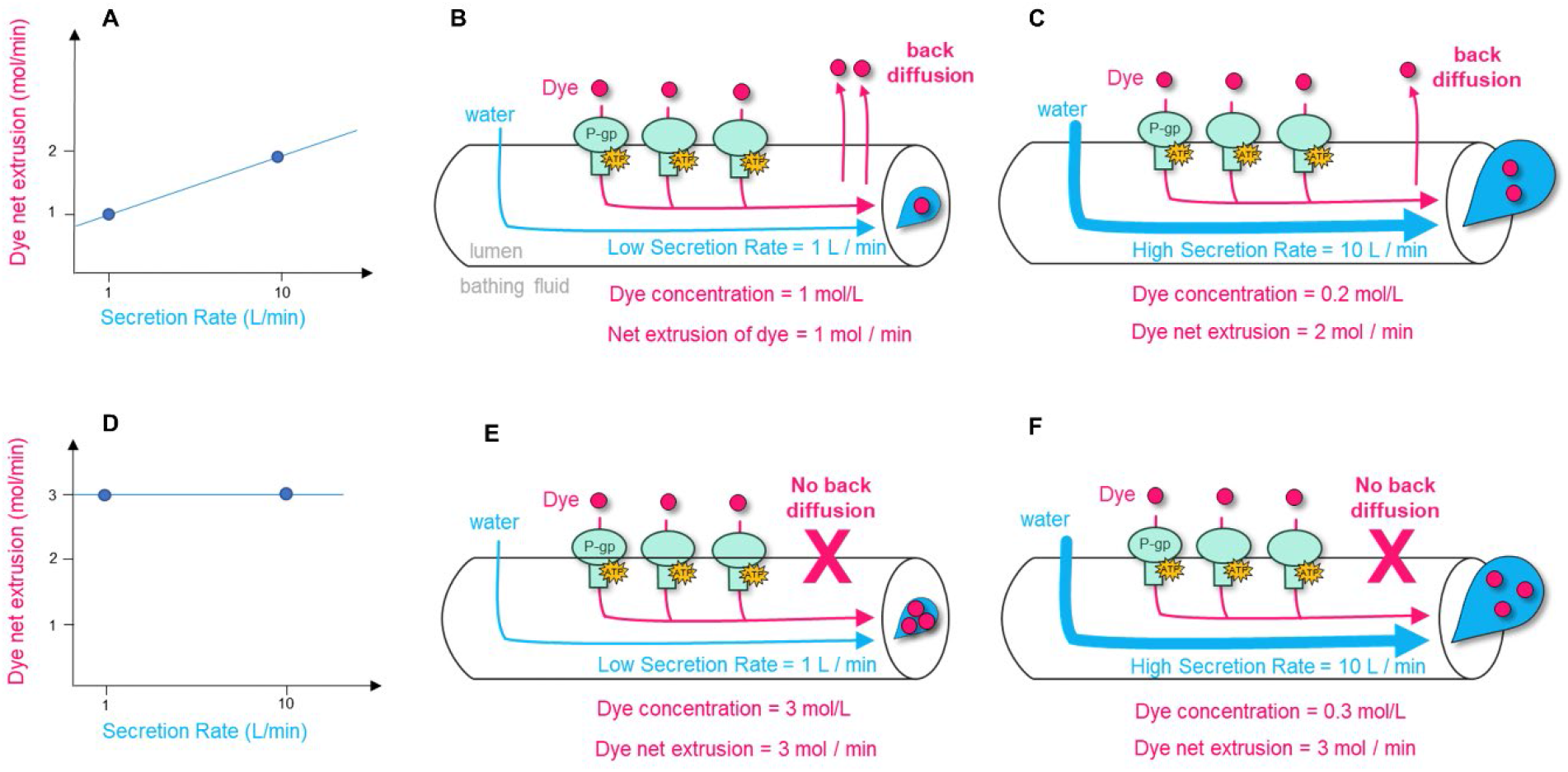
Schematic representation of the influence of the back diffusion on the net extrusion of a dye. (A) The net extrusion of a dye positively correlates with the fluid secretion rate when the dye can back diffuse from the lumen to the bathing fluid. (B) Indeed, at low fluid secretion rate, the dye concentration in the lumen rises, increasing the back diffusion and reducing the net transport of the dye. (C) Instead, at higher fluid secretion rates, the dye concentration in the lumen is diluted and the back diffusion is reduced, increasing the net transport of the dye. (D) If no back diffusion occurs, there is no correlation between the net transport of the dye and the fluid secretion rate. (E,F) At any rate of fluid secretion, the net transport of the dye remains constant, independently of the dye concentration in the lumen.

**S1 Table.**
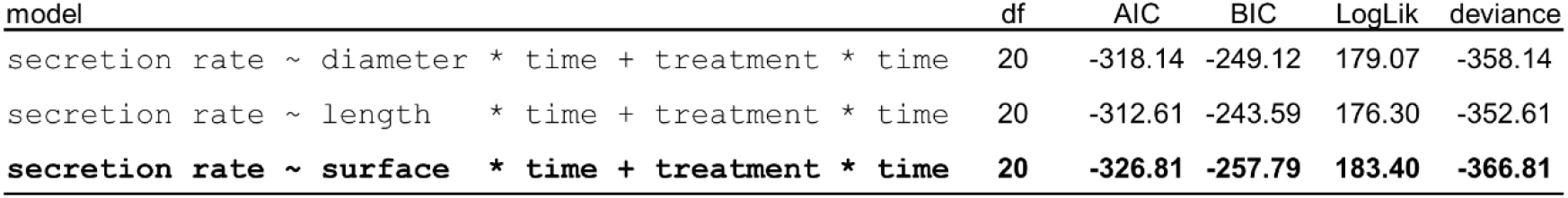
Comparison of linear mixed effect models incorporating length, diameter or surface area. Based on the lowest AIC parameter, the surface area was the best explanatory variable for the secretion rate. The row in bold indicates the model with the lowest AIC. Only the fixed effects are shown.

**S2 Table.**
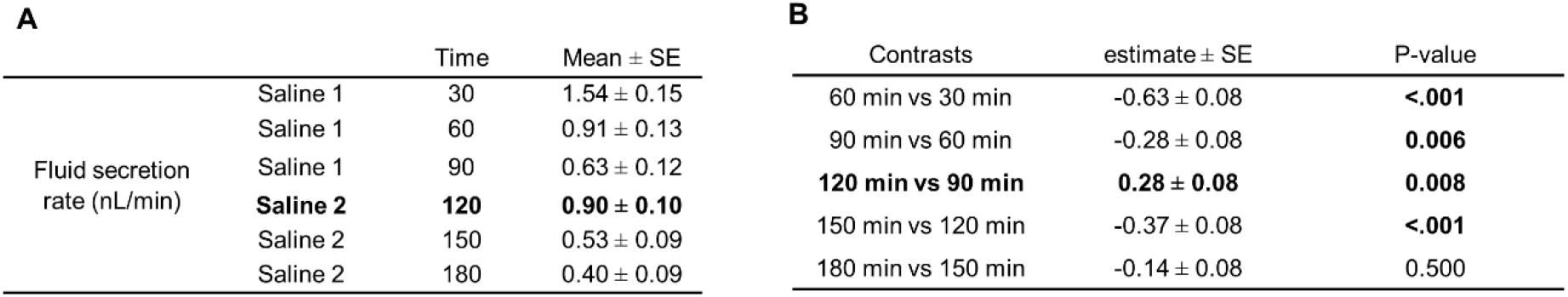
Outcomes of the linear mixed effect model investigating the effect the replacement of the saline with a fresh saline after 90 minutes of incubation upon the fluid secretion rate. The model applied was (Secretion rate ~ surface + time + (1| locust) + (1+time|tubule)). The rows in bold indicate the first observation after the saline has been replaced. (A) Summary of the mean values of fluid secretion rate at each time point. (B) Pairwise comparisons between subsequent times of incubation.

| **A** |  |  |  | **B** |  |  |
| --- | --- | --- | --- | --- | --- | --- |
|  |  | Time | Mean ± SE | Contrasts | estimate ± SE | P-value |
| Fluid secretion rate (nL/min) | Saline 1 | 30 | 1.54 ± 0.15 | 60 min vs 30 min | -0.63 ± 0.08 | **<.001** |
|  | Saline 1 | 60 | 0.91 ± 0.13 | 90 min vs 60 min | -0.28 ± 0.08 | **0.006** |
|  | Saline 1 | 90 | 0.63 ± 0.12 | **120 min vs 90 min** | **0.28 ± 0.08** | **0.008** |
|  | **Saline 2** | **120** | **0.90 ± 0.10** | 150 min vs 120 min | -0.37 ± 0.08 | **<.001** |
|  | Saline 2 | 150 | 0.53 ± 0.09 | 180 min vs 150 min | -0.14 ± 0.08 | 0.500 |
|  | Saline 2 | 180 | 0.40 ± 0.09 |  |  |  |

**S3 Table.** Outcomes of the linear mixed effect model investigating the diameter of the tubules in different moments during the Ramsay assay. The model applied was (diameter ~ assay time + (1| locust) + (1+time|tubule)). (A) Summary of the mean diameter of Malpighian tubules in vivo, at the beginning of the essay and at the end of the essay after 180 minutes of incubation. (B) Pairwise comparisons between different moments of the assay.

|  |  | Mean ± SE |
| --- | --- | --- |
| Diameter (µm) | In vivo | 71.0 ± 1.6 |
|  | Beginning assay | 69.3 ± 1.6 |
|  | End assay | 70.4 ± 1.6 |

**B**
|  | estimate ± SE | P-value |
| --- | --- | --- |
| Beginning assay vs in vivo | -1.7 ± 1.1 | 0.260 |
| end assay vs in vivo | -0.6 ± 1.1 | 0.847 |
| end assay vs beginning assay | 1.1 ± 1.1 | 0.551 |

